# Insertion and deletion evolution reflects antibiotics selection pressure in a *Mycobacterium tuberculosis* outbreak

**DOI:** 10.1101/2020.01.28.922765

**Authors:** Maxime Godfroid, Tal Dagan, Matthias Merker, Thomas A. Kohl, Roland Diel, Florian P. Maurer, Stefan Niemann, Anne Kupczok

## Abstract

In genome evolution, genetic variants are the source of diversity, which natural selection acts upon. Treatment of human tuberculosis (TB) induces a strong selection pressure for the emergence of antibiotic resistance in the infecting *Mycobacterium tuberculosis* (MTB) strains. MTB evolution in response to treatment has been intensively studied and mainly attributed to point substitutions. However, the contribution of insertions and deletions (indels) to MTB genome evolution remains poorly understood. Here, we analyzed a multi-drug resistant MTB outbreak for the presence of high-quality indels and substitutions. We find that indels are significantly enriched in genes conferring antibiotic resistance. Furthermore, we show that indels are inherited during the outbreak and follow a molecular clock with an evolutionary rate of 5.37e-9 indels/site/year, which is 23x lower compared to the substitution rate. Inherited indels may co-occur with substitutions in genes along related biological pathways; examples are iron storage and resistance to second-line antibiotics. This suggests that epistatic interactions between indels and substitutions affect antibiotic resistance and compensatory evolution in MTB.

**Author summary:** *Mycobacterium tuberculosis* (MTB) is a human pathogen causing millions of deaths every year. Its genome evolution has been intensively characterized through point substitutions, i.e., nucleotide exchanges that are inherited. Additional mutations are short or long insertions and deletions of nucleotides, termed indels. Short indels in genes might change the reading frame and disrupt the gene product. Here we show that antibiotic treatment has a strong impact on indel evolution in an MTB outbreak. Namely, indels occur frequently in genes causing antibiotic resistance upon disruption. Furthermore, we show that the molecular clock, i.e., the temporal emergence of variants over time, holds for short indels in MTB genomes. Finally, we observe that indels may co-occur with substitutions in genes along related biological pathways. These results support the notion that indels are important contributors to MTB evolution. We anticipate that including indels in the analyses of MTB outbreaks will improve our understanding of antibiotic resistance evolution.

## Introduction

*Mycobacterium tuberculosis* complex (MTBC) strains, the causative agents of tuberculosis (TB), are strict host-associated pathogens (1). With estimated numbers of ten million new infections and 1.2 million deaths in 2018 (2), TB is a major cause of human disease and mortality. In addition, *Mycobacterium tuberculosis sensu stricto* (MTB), the human-adapted member of the MTBC, has a high level of intrinsic and evolved antibiotic resistance (ABR), including multi-drug resistance (3). MTB genomes have a low genetic diversity and furthermore, comparative genomics of MTB genomes showed that genetic variation is only vertically inherited, likely due to the absence of horizontal transfer mechanisms in MTB (4,5). Consequently, MTB antibiotic resistance is considered to evolve *de novo* via point and segmental mutations and not by horizontal transfer of genetic material (6). Antibiotic resistance may induce high fitness costs that are frequently ameliorated by compensatory mutations (7). For example in MTB, mutations in *rpoB*, encoding the beta-subunit of the RNA polymerase, can lead to rifampicin resistance (8) and mutations in *rpoC* often compensate ABR-conferring mutations in *rpoB* (9,10). Notably, in asexual organisms, beneficial alleles are linked to the genetic background where they appeared. This results in competition between beneficial alleles (also known as clonal interference) and the hitchhiking of neutral or slightly deleterious alleles with beneficial ones. Indeed, time series patient sampling revealed that clonal interference and hitchhiking contribute to antibiotic resistance evolution in MTB (11,12).

Genetic variation in MTB strains is generally characterized by the emergence of substitutions that are observed as single-nucleotide polymorphisms (SNPs). Substitutions are the major source of variation in MTB genomes followed by insertions and deletions (indels). Short indels (up to 50 bp) were found to occur primarily in non-coding regions, in the repeat-containing PE-PPE genes and in ABR-conferring genes (13). Additionally, long insertions in MTB are mainly due to integration of the mobile element IS6110, a transposase-mediated insertion sequence (14). Importantly, previous studies analyzing MTB strain genome evolution provided evidence for the role of indels in ABR evolution (15–17).

Similarly to resistance determination, transmission dynamics within MTB outbreaks is generally inferred by SNP-based phylogeny reconstruction, after detecting SNPs from short-read sequencing data aligned to the complete and well characterized reference genome H37Rv (18,19). Outbreak reconstructions have furthermore been used to identify signals of positive selection in MTB strain evolution, for example, by identifying convergent evolution, i.e., variants that evolved independently multiple times. Convergent evolution in MTB has been observed in ABR-conferring genes (20) or in virulence factors (21). Furthermore, time-series sampling of MTB strains showed that substitutions in MTB genomes evolve at an approximately constant pace, i.e., substitutions follow a molecular clock (22). Notably, the substitution rate of MTB is on the lower end spectrum of prokaryotic substitution rates (6). Despite the low evolutionary rate observed for MTB strains, molecular dating can be used to infer the time of emergence of ABR-conferring substitutions (23) or the introduction time of strains into specific parts of the world (24).

Although previous studies extensively investigated the rate and impact of substitutions in MTB strain evolution, the contribution of indels has been sparsely analyzed. To address this question, we estimated the evolutionary rate and the phenotypic impact of insertions and deletions in MTB outbreak strains. In asexual organisms, genetic linkage is strong and might lead to epistatic interactions between variants. Hence, we investigated phylogenetically co-occurring indels and substitutions to describe their putative combined effects on MTB phenotype. As a paramount example for drug resistance evolution, we analyze a previously described multi-drug resistant clade of MTB lineage 2 (Beijing) strains, i.e., the Central Asian outbreak (CAO) (25). We further compare some aspects of indel evolution to the drug-susceptible lineage 4 (Euro-American) ‘Hamburg outbreak’ (26).

## Results

To study the evolution of point and segmental mutations in *M. tuberculosis*, we analyzed 353 MTB strains of the lineage 2 CAO that was detected previously to be involved in transmission of multi-drug resistant TB mainly in central Asia (Table S1a). Genetic variants were inferred by comparing the sample genomes to a closely related reference genome (strain *M. tuberculosis* 49-02 (25)). To assess the robustness of the genetic variation inference, we developed a back-genotyping approach, in which the inference procedure is performed against a simulated reference genome that includes the detected variants (Fig S1). Variants that are not supported by back-genotyping were considered as uncertain in this sample and variants that were inferred to be uncertain in many samples are unreliable and discarded from the analysis (Fig S2). Using our approach, we inferred in total 1806 high-quality variants in the CAO strains. These variants comprise a majority of SNPs (1598, 88.5%) and 208 insertions and deletions, where the majority of inferred indels are short (≤50bp; Table 1, Fig S3). We noted a peak in the distribution of insertion length around 1360bp that corresponds to 38 different insertions of the mobile element IS6110 (Fig S3). IS6110 insertions are known to be found preferentially in some genomic regions, i.e., insertional hotspots, where IS6110 insertions confer a growth advantage (14). The distribution of distances between IS6110 insertions in the CAO revealed seven insertional hotspots in the MTB genome, of which two hotspots have been previously described (Fig S4) (14).

**Table 1.** Summary and genomic localization of detected variants.

|  | SNPs | Insertions |  | Deletions |  | Total |
| --- | --- | --- | --- | --- | --- | --- |
|  |  | Short | Long | Short | Long |  |
| **In gene** | 1362 (85.23%) | 48 (73.85%) | 28 (63.64%) | 65 (73.86%) | 11 (100%) | **1514** |
| **Intergenic** | 202 (12.64%) | 13 (20%) | 13 (29.54%) | 19 (21.59%) | - | **247** |
| **In pseudogene** | 34 (2.13%) | 4 (6.15%) | 3 (6.82%) | 4 (4.55%) | - | **45** |
| **Total** | **1598** | **65** | **44** | **88** | **11** | **1806** |
| **Parsimony informative** | 434 (27.2%) | 18 (27.7%) | 14 (31.8%) | 16 (18.2%) | 4 (36.4%) | **486** |
| **Compatible** | 394 (90.8%) | 11 (61.1%) | 10 (71.4%) | 13 (81.2%) | 2 (50%) | **428** |

Percentages are calculated based on the total number of variants in each variant class, except for the compatible variants, where the percentage is calculated based on the number of parsimony informative variants in each variant class. The majority of SNPs and indels are found in coding regions. The distribution of variants in protein-coding and intergenic regions differs significantly from the expectation by chance (Fisher's exact test, $p < 0.01$ ), where 247 (13.7%) variants are inferred in intergenic regions that span 7.8% of the genome.

### Short indels contribute significantly to antibiotic resistance evolution

To infer the putative phenotypic impact of the inferred variants in coding regions we examined their localization in genes of known function. For this purpose, we retrieved a list of ABR-conferring genes (Table S2), i.e., genes where mutations were found to confer antibiotic resistance. Additionally, we classified the MTB genes in two categories of essentiality, according to their requirement for growth *in vitro* (i.e., essential) or not (i.e., dispensable) (27). In particular, we investigated the distribution of genetic variants in essential and ABR-conferring genes. Depletion of variants in specific gene categories indicates purifying selection acting on that category, whereas enrichment serves as an indication for positive selection.

First, we observed a fourfold depletion of short indel frequency in essential genes; the distribution of SNPs, however, is not significantly different between essential and dispensable genes (Fig 1a). Furthermore, SNPs and short indels are enriched in ABR-conferring genes compared to the remaining genes (Fig 1b). When we classify the ABR-conferring genes into essential (27, 29.4% of ABR-conferring genes) and dispensable (65, 70.6% of ABR-conferring genes), we observed that SNPs are enriched both in ABR-conferring genes that are essential and in ABR-conferring genes that are dispensable. In contrast, short indels are significantly enriched in the ABR-conferring genes that are dispensable but not in the essential ABR-conferring genes (Fig 1c,d). In comparison, the drug-susceptible Hamburg outbreak did not show variants in ABR-conferring genes (Fig S5).

**Fig 1.**
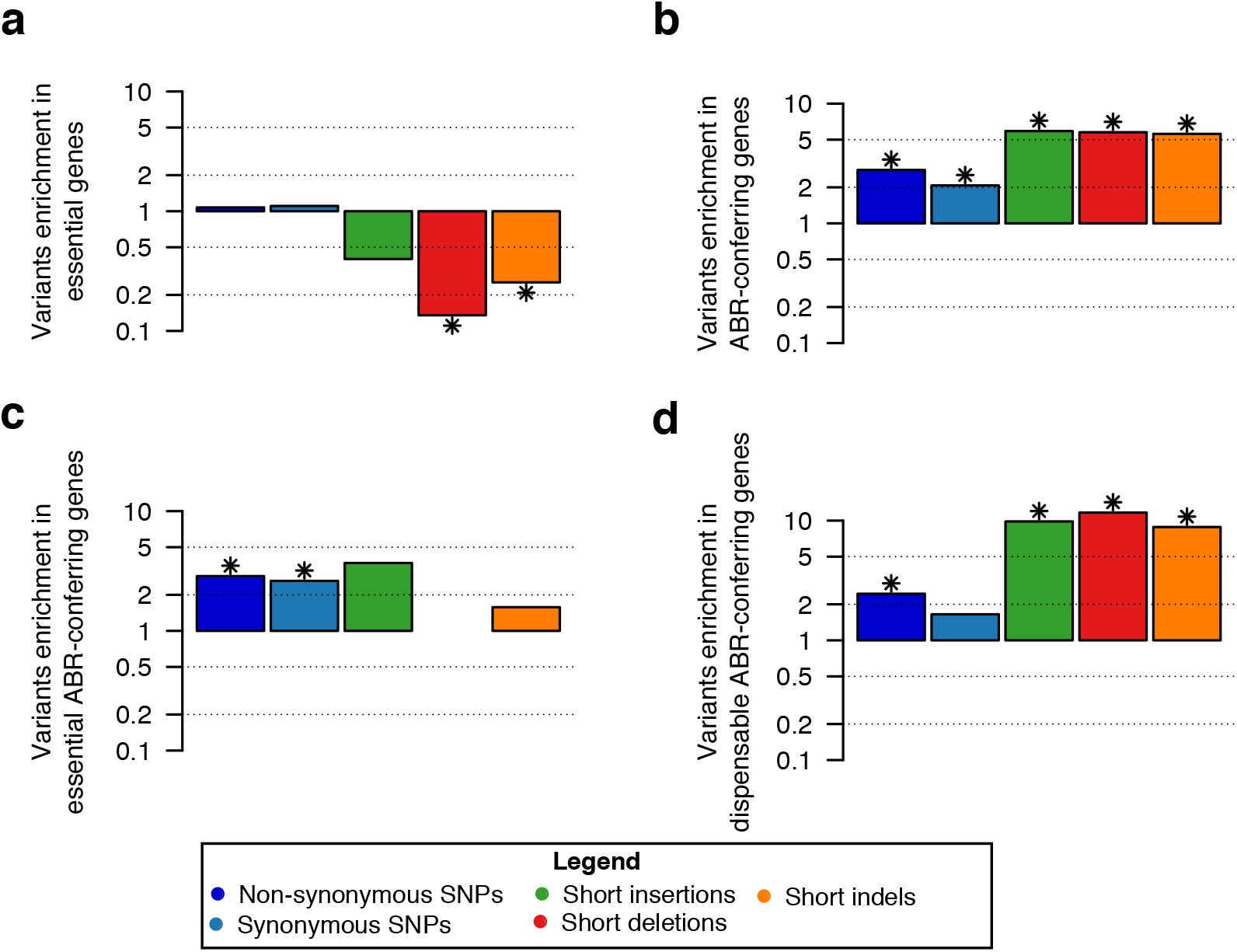
Enrichment analyses of variants in gene categories. The variants enrichment is calculated as the ratio of the proportion of genes with variants in a gene category and the proportion of genes with variants outside the gene category. For example, three essential genes (0.6% of all essential genes) have short indels and the remaining short indels occur in 94 dispensable genes (3.3% of all dispensable genes), which results in a variants ratio of short indels in essential genes of 0.25, i.e., a fourfold depletion. We show the ratio for the gene categories (a) Essential, (b) ABR-conferring, (c) Essential and ABR-conferring, and (d) Dispensable and ABR-conferring. Significant enrichment or depletion, marked by a star, is estimated using Fisher’s exact test (p-value < 0.05, corrected for FDR, Table S3).

The enrichment analyses highlight the selection pressures on SNPs and indels in the CAO. The depletion of short indels in essential genes provides evidence for the presence of strong purifying selection against indels in essential genes. In addition, the observed enrichment in ABR-conferring genes likely stems from the strong selection pressure on antibiotic resistance in the multi-drug resistant CAO. The significant enrichment of short indels in ABR-conferring genes that are dispensable shows that indels contribute to the evolution of antibiotic resistance in a highly resistant outbreak, potentially by frameshifts that disrupt the protein sequences.

### Insertions and deletions contribute phylogenetic signal in an MTB outbreak

To study the transmission of indels in an outbreak, we next describe how the genetic variants are inherited in the CAO. To this end, we reconstructed the outbreak phylogeny from the presence-absence pattern of the variants in the strain genomes, where uncertain variants in a sample correspond to gapped positions. This analysis revealed that SNPs are the main contributors to the phylogenetic signal, where most of parsimony informative SNPs are compatible with the phylogeny (Table 1). The inclusion of indels in the phylogenetic reconstruction increases the resolution of the tree topology at multiple places, where six internal branches are supported by a single short indel only (Fig 2). In comparison, the analysis of 64 samples of the ‘Hamburg outbreak’ resulted in 112 variants, of which the majority are SNPs as well (Fig S5a). In the Hamburg outbreak phylogeny, two internal branches are supported by indels only (Fig S5b).

**Fig 2.**
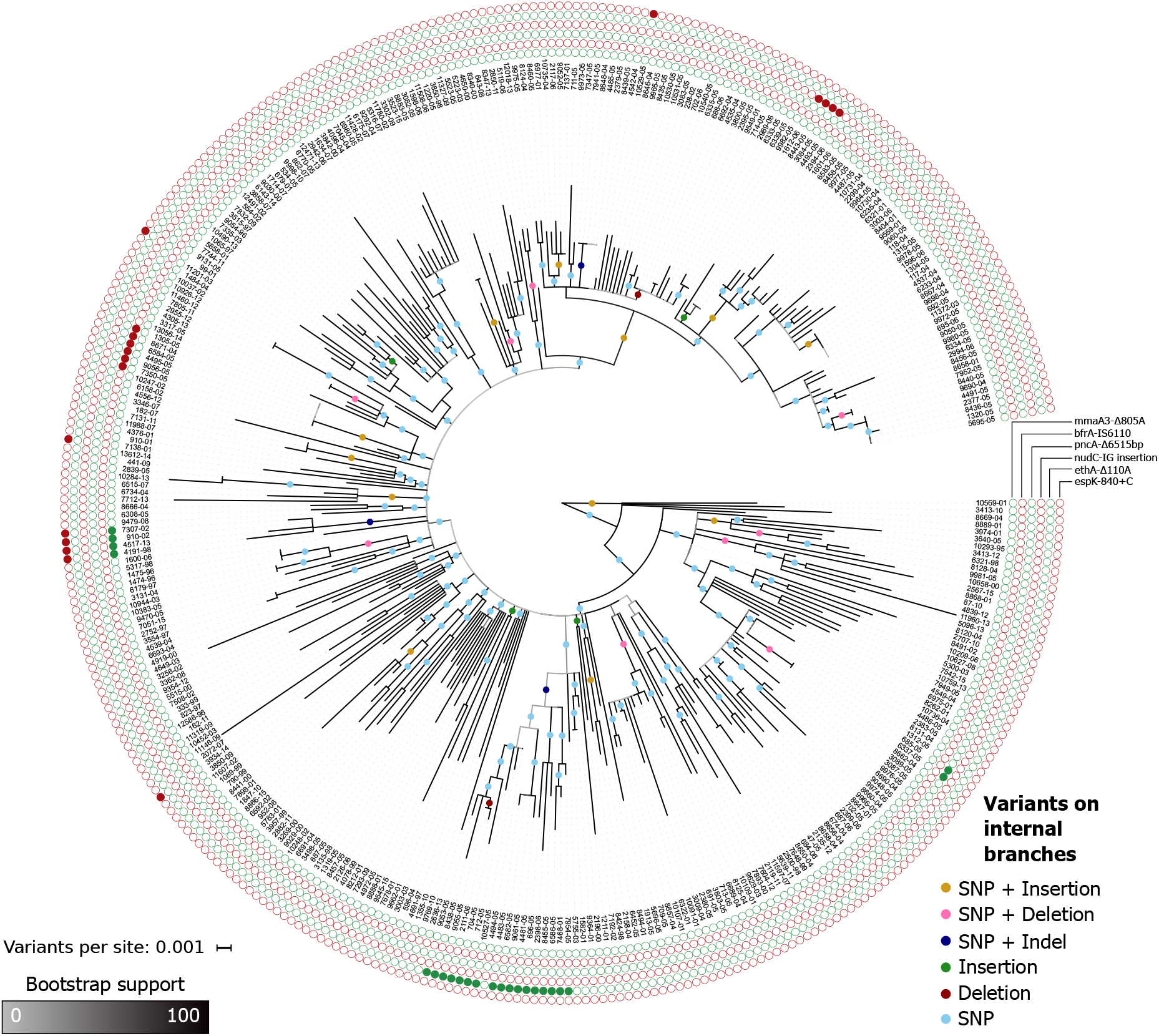
Phylogenetic tree of the CAO. 486 (26.9%) of the variants are parsimony informative, and 56 variants (3.1%) are incompatible with the tree topology (Table 1). The root position is the temporal root estimated by dating the phylogeny with LSD. Circles on branches represent variants that are compatible with the branch, i.e., they likely have emerged on that branch. We found that six branches in the tree have only short indels (four branches with insertions, two branches with deletions). The outer circles show example variants that are highlighted in the text.

We also compared our approach to the standard SNP-based phylogenetic approach for the CAO data. The complete tree contains 157 internal branches of length larger than zero, whereas the SNP phylogeny contains 149 internal branches of length larger than zero. Notably, 138 branches (92.6% of the branches in the SNP phylogeny) are found in both trees. Thus, our phylogenetic inference is consistent with the standard SNP-based phylogeny, where the resolution of additional branches indicates that indels provide additional insights into possible transmission events.

### Substitutions and short indels in the CAO follow a molecular clock

To compare the pace of substitutions and indels in the outbreaks, we examined their evolutionary rates on the phylogeny. We found that substitutions passed the stringent test for temporal signal, with an estimated rate of 1.23e-7 substitutions/site/year (Fig 3). For comparison, substitutions in the Hamburg outbreak passed the intermediate test for temporal signal, with an estimated rate of 7.51e-8 substitutions/site/year (Fig S5c). This rate is lower compared to the CAO rate; however, the confidence intervals of the estimates overlap (Fig 3, S5c). The substitution rate estimated here for the CAO is within previously estimated rates for lineage 2 (22). Furthermore, there is a known difference in the rate of evolution between MTB lineage 2 and lineage 4 (28), which is consistent with our estimates for the lineage 2 CAO and the lineage 4 Hamburg outbreak.

**Fig 3.**
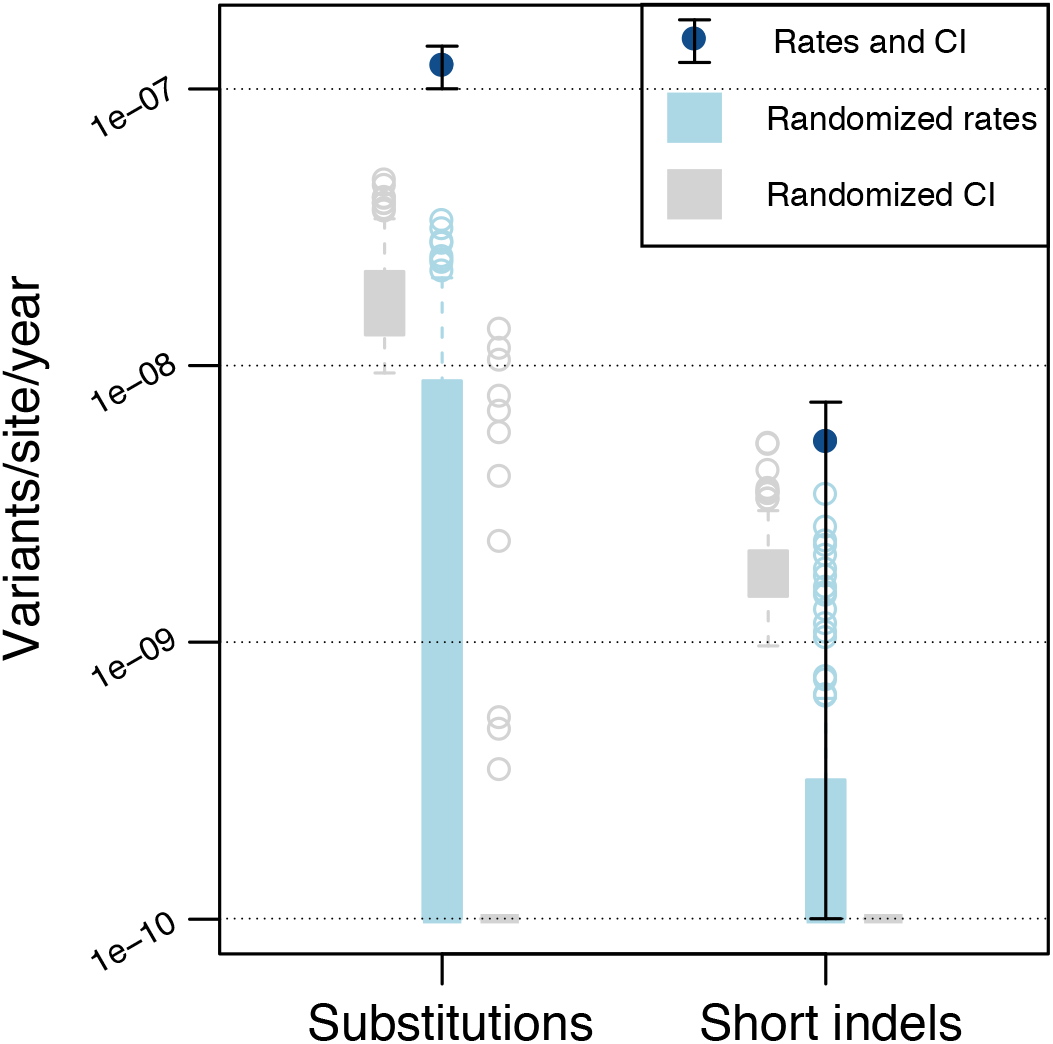
Evolutionary rates for substitutions and short indels and their associated 95% confidence intervals (CI) estimated with LSD. Substitution rate is estimated at 1.23e-7 [1.00e-7–1.43e-7] substitutions/site/year and the short indel rate is estimated at 5.37e-9 [1e-10–7.37e-9] indels/site/year. The randomized rates are estimated with the date-randomization test (29). For substitutions, there is no overlap in rate and confidence interval between real data and randomized data, indicating temporal signal accorded by the stringent test. For short indels, the rate of the real data does not overlap with the rates of the randomized data; however, the confidence interval of the real data overlaps with the confidence intervals of the randomized data, which indicates temporal signal of short indels accorded by the simple test, is weaker than that of substitutions. Due to the limited number of events, there is not sufficient temporal signal to estimate evolutionary rates for short insertions and deletions separately or for long indels.

We then estimated the evolutionary rate of indels in the CAO. Short insertions and deletions are assumed to emerge by similar point mutation processes, in contrast to long indels that are due to segmental mutations (e.g., (30)). We thus considered short indels and long indels separately for the rate estimation, where only short indels have temporal signal (according to the simple test), with a rate of 5.37e-9 short indels/site/year (Fig 3). Hence, our analysis revealed that substitutions and short indels follow a molecular clock and that short indels evolve 23 times slower than substitutions in MTB lineage 2 strains. The difference between substitution and indel rates might be explained by the extremely efficient and redundant MTB repair mechanisms. Error-prone repair mechanisms, such as the DnaE2 pathway that is involved in trans-lesion synthesis, are known to introduce substitutions (31). Thus, MTB repair mechanisms might generate a mutational bias towards substitutions, leading to a stable genome with few structural variations over evolutionary time.

### Indels are subject to vertical inheritance and convergent evolution in the CAO

To describe the function of variants that are transmitted in the outbreak or that evolved multiple times independently, we explored the phylogenetic distribution and congruence of indel variants in the reconstructed CAO tree. Parsimony informative variants that are compatible with the phylogeny (termed compatible variants hereafter) are inferred as vertically inherited and transmitted to multiple hosts; thus, their effect on MTB fitness is likely not deleterious or even advantageous. In addition, incompatible variants are convergent events, where the same variant occurs independently in two (or more) disparate branches of the tree, indicating convergent evolution. Convergent variants might serve as evidence for positive selection if they emerge in similar genetic backgrounds and have identical phenotypic impact (32).

We found 36 compatible and 16 incompatible indels (31.8%), where incompatible indels are enriched among the parsimony informative indels in comparison to incompatible SNPs (40, 9.22% of all parsimony informative SNPs, p-value<0.01, Fisher’s exact test). This difference can be traced back to convergent short indels in homopolymer regions and also inference bias (Table S4). We found that 30 of the 36 compatible indels are located in coding regions (83.3%; Table S4), out of which 29 are in dispensable genes and two are in ABR-conferring genes (*mmaA3* and *ethA*; Fig 2). In contrast, nine of the 16 incompatible indels are found in coding regions (56.2%; Table S4). Thereof, eight indels are found in dispensable genes and two in ABR-conferring genes, where the only incompatible indels in ABR-conferring genes are long deletions completely or partially deleting *pncA*. In contrast, incompatible SNPs are mainly found in genes conferring antibiotic resistance (21 of 29 incompatible SNPs in coding regions, 72.4%, Table S5). This is in agreement with previous results where convergent SNPs have been observed in MTB genes that confer antibiotic resistance and compensatory mechanisms (17,20).

We next considered genes in which multiple indels were inferred, i.e., genes affected by convergent evolution due to indels. We found 15 indels affecting four ABR-conferring genes; in three of these genes (*rpoB*, *tlyA*, *ethA*) additional SNPs were inferred in different samples, whereas no SNPs were inferred in *pncA* (Table S6). In addition, twelve genes where multiple indels have been found do not confer antibiotic resistance (e.g., *espB* and four PE genes; Table S6); these genes do not contain SNPs in any sample.

PE and PPE are repeat-containing genes that are secreted and they are hypothesized to be important for MTB interaction with the host immune system (33). An examination of all variants inferred in PE and PPE genes in the CAO strains revealed 17 short indels. Of these, only three (17.6%) cause a frameshift, which is much lower than the proportion of frameshift-causing indels in all coding sequences (77.8%). Furthermore, we found seven parsimony informative short indels, out of which six are compatible (five in-frame) and one is an incompatible in-frame deletion. The vertical inheritance and the enrichment of in-frame indels in PE and PPE genes indicate that these proteins are fast evolving, further supporting the hypothesis that they are involved in host recognition (33).

Taken together, we found 52 parsimony informative indels in 34 different genes. It is remarkable that only two of these genes (*ethA* and a nitronate monooxygenase) likely evolved under positive selection as inferred by the ratio of nonsynonymous to synonymous substitutions (dN/dS>1, Table S7). We note that the inference of positive selection can only be performed for three of the 34 genes with parsimony informative indels due to the lack of SNPs in the remaining genes. We further discovered that four genes with parsimony informative indels (11.8%) are included in a set of 116 (3%) genes that were found to be under positive selection in a recent survey of dN/dS in MTB (34). Thus, while indels can be found in genes that are under positive selection as calculated by the dN/dS ratio, they might also uncover additional genes involved in adaptation. In the following, we study six example indels in detail that were selected to highlight convergent evolution, inherited antibiotic resistance, and putative epistatic interactions.

#### A short deletion shortens the ABR-conferring gene *mmaA3*

A compatible deletion of one base pair was observed in the gene *mmaA3* in four related samples (Fig 2). The deletion results in a frameshift and a premature stop codon yielding a truncated protein sequence (Fig 4a). The protein MmaA3 acts along the synthesis pathway of mycolic acids, which are essential components of the bacterial membrane (35). The gene is classified as ABR-conferring, yet it is classified as dispensable *in vitro*. In addition, we observed five SNPs and one IS6110 insertion that co-occur with the 1bp deletion in the same four samples (Table 2). Three of the five SNPs are non-synonymous substitutions in genes that encode proteins involved in membrane biogenesis (Table 2). Our results thus revealed several substitutions and indels, which emerged and were vertically inherited together, and which likely have an effect on the function of membrane biosynthesis genes.

**Table 2.** Variants that co-occur with the example indels (Fig 2).

| Genes with focus variant | Co-occurring variant | Locus tag | H37Rv homolog | Gene name | Product | Notes |
| --- | --- | --- | --- | --- | --- | --- |
| *mmaA3* | sSNP | MT49_RS03380 | Rv0645c | *mmaA1* | mycolic acid methyltransferase MmaA1 | Involved in membrane biogenesis |
| *mmaA3* | Intergenic SNP | MT49_RS08155 | Rv1535 |  | hypothetical protein |  |
| *mmaA3* | IS6110 | MT49_RS16445 | Rv3126c |  | hypothetical protein |  |
| *mmaA3* | nsSNP | MT49_RS17845 | Rv3383c | *idsB* | polyprenyl synthetase family protein | Involved in membrane biogenesis |
| *mmaA3* | nsSNP | MT49_RS19685 | Rv3740c |  | wax ester/triacylglycerol synthase family O-acyltransferase | Involved in membrane biogenesis |
| *mmaA3* | nsSNP | MT49_RS20030 | Rv3800c | *pks13* | acyltransferase domain-containing protein | Involved in membrane biogenesis |
| *bfrA* | nsSNP | MT49_RS01400 | Rv0265c |  | ABC transporter substrate-binding protein | Probable periplasmic iron-transport lipoprotein (mycobrowser) |
| *bfrA* | nsSNP | MT49_RS03540 | Rv0676c | *mmpL5* | siderophore RND transporter MmpL5 | ABR-conferring |
| *bfrA* | sSNP | MT49_RS04920 | Rv0935 | *pstC1* | phosphate ABC transporter permease |  |
| *bfrA* | nsSNP | MT49_RS08525 | Rv1625c | *cya* | adenylate cyclase |  |
| *bfrA* | nsSNP | MT49_RS09530 | Rv1830 |  | MerR family transcriptional regulator |  |
| *bfrA* | sSNP | MT49_RS09595 | Rv1843c | *guaB1* | GuaB1 family IMP dehydrogenase-related protein |  |
| *bfrA* | nsSNP | MT49_RS12315 | Rv2337c |  | hypothetical protein |  |
| *bfrA* | nsSNP | MT49_RS20315 | Rv3854c | *ethA* | FAD-containing monooxygenase EthA | ABR-conferring |
| *pncA* | Intergenic SNP | MT49_RS03955 | Rv0755c | PPE12 | PPE family protein PPE12 |  |
| *pncA* | sSNP | MT49_RS13630 | Rv2582 | *ppiB* | peptidyl-prolyl cis-trans isomerase |  |
| *ethA* | nsSNP | MT49_RS03490 | Rv0667 | *rpoB* | DNA-directed RNA polymerase subunit beta | Essential ABR-conferring gene |
| *ethA* | Intergenic IS6110 | MT49_RS09400 | Rv1804c |  | hypothetical protein | predicted secreted protein |

Gene names are displayed according to the Mycobrowser annotations (https://mycobrowser.epfl.ch/, last accessed 28 November 2019), when available. Locus tags and products are taken from the annotation of the 49-02 reference genome. Notes show additional information related to protein functions.

**Fig 4.**
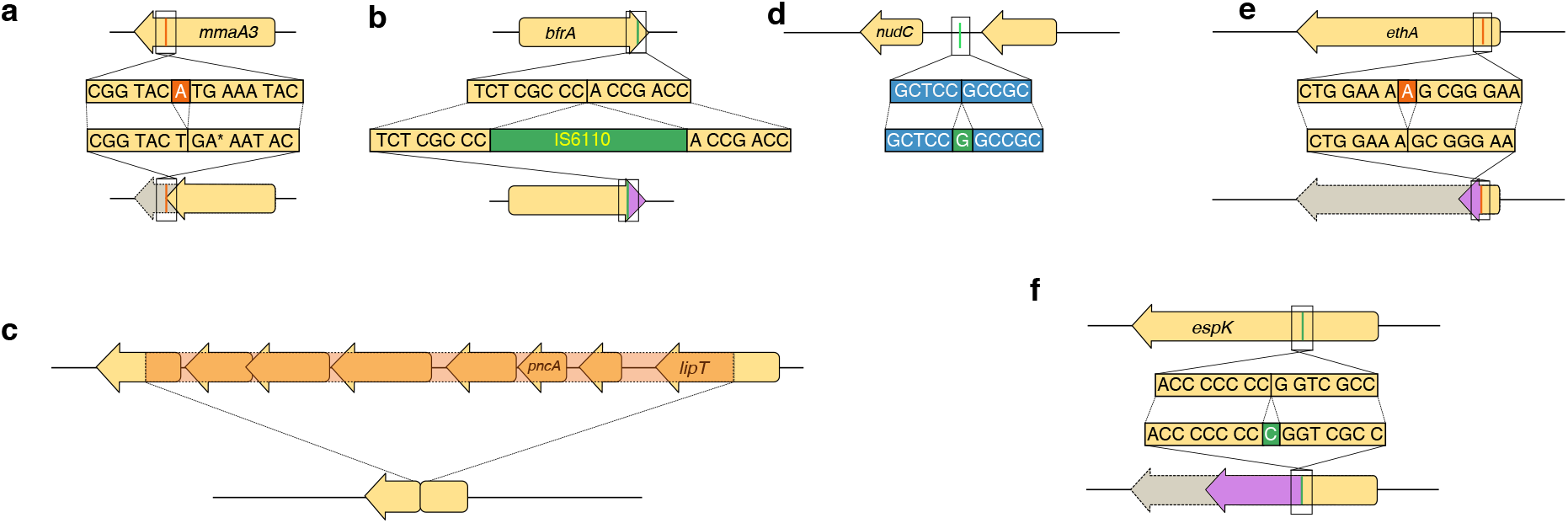
Examples of indels subject to vertical inheritance and convergent evolution. The orientation of the gene is relative to the reference genome 49-02. At the top is the ancestral (reference) sequence and at the bottom the evolved sequence. See Fig 2 for phylogenetic locations. Gene names are displayed according to the Mycobrowser annotations (https://mycobrowser.epfl.ch/, last accessed 28 November 2019), when available. Following annotations are shown as (Locus tag in 49-02, homolog in H37Rv). (a) Single base-pair deletion close to the 5’ end of *mmaA3* (MT49_RS03370, Rv0643c). The deletion removes the residue at position 805 of the coding sequence, resulting in a stop codon where the mutated protein (269 amino acids) corresponds to 91.5% of the wild-type protein. Six variants co-occur on the same internal branch (Table 2). (b) IS6110 integration close to the 3’ end of the *bfrA* gene (MT49_RS09760, Rv1876). The integration occurs at position 471 of the coding sequence (98.1% of total CDS length), resulting in a protein of 162 amino acids (2 amino acids longer than the wild-type), with the last two amino acids of the wild type different in the mutated protein. We find eight co-occurring variants on the same internal branch (Table 2). (c) 6515 base-pair deletion removing the *pncA* gene (MT49_RS10715, Rv2043c) and five of its neighboring genes. The left breakpoint is located at position 529 in the gene *ugpC* (MT49_RS10690, Rv2038c). The right breakpoint is located at position 453 in a gene encoding for a carboxylesterase/lipase family protein (MT49_RS10725, Rv2045c). Four of the deleted neighboring genes encode for ABC transporters (Table S4). Two variants co-occur with this long deletion (Table 2). (d) Intergenic single base-pair insertion 39bp upstream of the *nudC* gene (MT49_RS16870, Rv3199c). We could not detect the promoter sequence in this intergenic region using BPROM (36). (e) Single base-pair deletion in the beginning of the *ethA* gene (MT49_RS20315, Rv3854c). This deletion occurs at position 110 of the coding sequence (7.5% of the CDS length), which results in a frameshift where the resulting protein is truncated with a length of 62 amino acids long (12.6% of the wild type length). This deletion co-occurs with two variants (Table 2). (f) Single base-pair insertion in the *espK* gene (MT49_RS20440, Rv3879c). This insertion occurs at position 840 of the coding sequence (37.7%), resulting in a truncated protein of 465 amino acids (62.7% of the wild type length).

#### An IS6110 insertion elongates a bacterioferritin gene

A compatible insertion of an IS6110 element was identified at the 3’ end of *bfrA* in six related samples (Fig 2; Fig 4b). The deleterious effect of indels at the 3’ end of genes is considered minimal due to the indel location at the end of the open reading frame, thus maintaining the majority of the coding sequence (37). Indeed, the IS element insertion yields a protein sequence that is two amino acids longer than the wildtype, where 158 amino acids (98.8%) of the wild type protein are retained (Fig 4b). BfrA is an essential component for iron storage and distribution depending on iron availability (38) and it is classified as dispensable *in vitro*. We observed eight additional SNPs that co-occur with the IS6110 insertion in the same six samples (Table 2). Two of the SNPs are non-synonymous substitutions in ABR-conferring genes: *mmpL5* and *ethA*. MmpL5 is annotated as a siderophore transporter and is associated with bedaquiline resistance; EthA is associated with resistance to ethionamide (3). Of the remaining co-occurring SNPs, one non-synonymous substitution is observed in a gene encoding for a protein related to iron export and one non-synonymous substitution is found in a MerR transporter family that plays a role in responding to environmental stresses, such as oxidative stress, heavy metals or antibiotics (39) (Table 2). Thus, the IS1660 insertion in *bfrA* might hitchhike with the co-occurring substitutions in *mmpL5* and *ethA* or it might even be a compensatory variant for these substitutions that confer antibiotic resistance and may be additionally related to iron transport and storage.

#### A long deletion completely removes the ABR-conferring *pncA* and neighboring genes

A compatible 6515bp deletion of *pncA* with five neighboring genes was observed in two related samples (Fig 2). This deletion furthermore disrupts two additional neighboring genes and results in a chimeric coding sequence (Fig 4c). PncA activates pyrazinamide, a first-line antibiotic of the category “prodrug”, i.e., a compound that needs to be activated to exhibit toxicity. The disruption of *pncA* renders pyrazinamide inactive and thus results in antibiotic resistance; hence, indels in *pncA* have been previously observed to confer resistance to pyrazinamide (40). In our data, in addition to the multi-gene deletion, we observed a complete *pncA* deletion and nine disruptive indels across the tree (four short insertions, one long insertion, and four long deletions). Thus, *pncA* has the highest frequency of convergent indels in the CAO.

#### An intergenic short insertion is located upstream of the ABR-conferring *nudC*

A compatible 1bp insertion located in an intergenic region 39bp upstream of *nudC* was observed in four related samples (Fig 2, Fig 4d). NudC is an NAD(+) diphosphatase, where antibiotic resistance to isoniazid and ethionamide was observed when overexpressing *nudC* (41). Of note, the *nudC* gene in the outbreak reference genome 49-02 has a 239P->239R polymorphism compared to the reference H37Rv. This position has been found to disrupt the NudC dimer formation, hence it is expected to affect the NudC catalytic activity. The observed indel is compatible; hence, we hypothesize that the short insertion is advantageous to MTB, for example by altering the *nudC* expression level. An increased *nudC* expression might confer antibiotic resistance to isoniazid and/or ethionamide.

#### A short deletion disrupts the ABR-conferring *ethA*

A compatible 1bp deletion was observed in the *ethA* gene in 17 samples with an additional uncertain sample (Fig 2). This deletion results in a frameshift leading to a truncated protein (Fig 4e). EthA, an FAD-containing monooxygenase, is involved in the activation of ethionamide, a second-line antibiotic prodrug. The downregulation of *ethA* has been demonstrated to generate an ethionamide resistance phenotype (42). We observed two additional co-occurring variants in the same samples, where one is a non-synonymous substitution in *rpoB* (Table 2). We hypothesize that the disruption of EthA confers a strong selective advantage by mediating resistance to ethionamide. RpoB is known to confer resistance to rifampicin and mutations outside the rifampicin resistance determining region might compensate the cost of the resistance (10). Here we observed a substitution outside the rifampicin resistance determining region and thus hypothesize that it might be involved in compensation, resulting in the vertical inheritance of the variants. Interestingly, we find 20 additional variants in *ethA* throughout the tree, of which 17 are non-synonymous SNPs and three are frameshift single base pair deletions. The presence of SNPs that are exclusively non-synonymous shows that this gene is under strong positive selection (Table S7).

#### An incompatible short insertion disrupts the type VII secretion system gene *espK*

We observed a 1bp insertion in the *espK* gene in eight samples including four related samples and four unrelated samples (Fig 2). Hence this insertion likely emerged five times independently, indicating convergent evolution. The variant emerged in a homopolymer region of seven cytosines (Fig 4f), resulting in a frameshift and a premature stop codon. The *espK* gene is located in the ESX-1 locus, a type VII secretion system. The locus additionally comprises PE and PPE genes, encoding for proteins that are exported or found in the cell membrane (43). EspK is thought to act as a chaperone of the neighboring *espB* gene (44), and is found dispensable for growth *in vitro*. EspB acts as a repressor of the host immune response, thereby increasing MTB survival (45). Notably, it was shown that inhibition of *espK* and *espB* results in reduced virulence in comparison to the wild-type (45). The 1bp insertion likely renders EspK nonfunctional and hence has a direct effect on EspB function as well. Previous studies observed convergent substitutions in another type VII secretion system gene (*esxW* in ESX-5) that increased MTB transmissibility (21). Hence, the 1bp indel in the type VII secretion system might be related to MTB transmissibility as well.

Our examples demonstrate that indels contribute to the evolution and diversification of MTB CAO strains, by affecting essential metabolic pathways and antibiotic resistance with potential pathobiological consequences. Furthermore, we found that compatible indels often co-occur with substitutions that affect related functions or pathways (Table 2). These co-occurrences could be explained by epistatic interactions in which either indels compensate the effects of substitutions or *vice versa*. In addition, the significant enrichment of short indels in ABR-conferring genes that are dispensable shows that indels contribute significantly to ABR evolution in the multi-drug resistant CAO, likely by frameshifts that disrupt the protein sequences. Our study demonstrates that, even if rare, including indels in outbreak genome analyses supplies crucial evidence for the profiling of antibiotic resistant strains and might reveal epistatic interactions.

## Discussion

The contribution of indels to genome evolution is often understudied, mainly due to difficulties in reliable indel detection methodology. Our approach allows to infer high-quality indels by estimating the level of inference uncertainty, which was used to identify unreliable genetic variants. Applying this approach to MTB sequencing data, we show that indels can be employed to increase the resolution of MTB strain comparisons in genomic epidemiology approaches, e.g., for outbreak investigations. The accurate detection of indels in the CAO revealed an indel evolutionary rate that is lower than the substitution rate. Finally, indels are an important factor in the evolution of antibiotic resistance in MTB, where compatible and convergent indels represent putative targets for positive selection and where co-occurring variants highlight epistatic interactions.

MTB outbreak reconstructions so far mainly relied on the estimation of phylogenies based on SNPs. Here we show that six branches of the CAO tree and two branches of the Hamburg outbreak tree were reconstructed solely by short indels. Furthermore, the comparison of the CAO tree inferred solely from SNPs and inferred from SNPs and indels revealed similar topologies. Thus, the inclusion of indels can refine outbreak phylogenies.

The MTB mutation rate, that is, the rate at which mutations arise in the genome, is in the range of bacterial mutation rates (between 1.4e-10 for *Thermus thermophilus* and 4e-9 for *Buchnera aphidocola*; 1.9e-10 mutations/bp/generation for *M. tuberculosis* (46)). However, MTB strain evolution is characterized by a long generation time (1) and strong purifying selection that eliminates most genetic variants from the population, with only few mutations being fixed (47). Both of these processes contribute to a low substitution rate in MTB strains compared to strains of other bacterial species (6). A previous comparison between evolutionary rates of mutations and indels in multiple bacterial species showed a 2.8 to 9.7-fold decrease of indel rates compared to mutation rates (48). Notably, the comparison in the latter study is based on *de novo* rates, i.e., variants that are arise in a bacterial individual per generation, which aim to include all variants before selection. In contrast, outbreak analyses include only the variants that are observed after the effect of selection. Hence, the 23-fold decrease of the MTB indel rate compared to the substitution rate can reflect both a lower *de novo* indel rate, as observed for other bacterial species, and a stronger effect of purifying selection on indels. The latter is expected since indels incur a higher fitness cost than substitutions as they often disrupt genes and render a truncated gene product (48).

Notably, MTB is evolving strictly vertically without the contribution of recombination. The advantage of sex and recombination is widely discussed (e.g., (49)). On the one hand, recombination is beneficial by combining advantageous alleles from different genotypes in the population; whereas in the absence of recombination, advantageous alleles are linked to the genetic background where they arise (the Hill-Robertson effect). This genetic linkage might result in the fixation of neutral or slightly deleterious alleles by genetic hitchhiking with advantageous alleles. In addition, clonal interference between beneficial alleles in different genetic backgrounds slows down adaptation (the Fisher-Muller effect). On the other hand, in the presence of positive epistasis, i.e., when the double mutant has a higher fitness than expected from the individual alleles, recombination can lead to a decrease in fitness by breaking up advantageous allele combinations (resulting in recombination load). It has been observed that the magnitude of recombination impacts the genetic architecture of a species, where positive epistasis evolved in a bacteriophage model system under low recombination but not under high recombination (50). In addition, an artificial gene network model has been used to demonstrate that positive epistasis evolves in asexual populations whereas negative epistasis can evolve in sexual populations (51). It is thus expected that the asexual lifestyle of MTB results in a genetic architecture with widespread positive epistasis.

Indeed, epistatic interactions between genetic variants are widespread in MTB, where the emergence of ABR-conferring mutations is often accompanied by compensatory mutations (3). The fixation of compensatory variants might even be favored over the reversal of antibiotic resistance in the absence of ongoing antibiotic treatment, because compensatory mutations might appear with a higher rate compared to the very specific target of a reversal mutation (52). After the compensatory mutation increased in frequency, a newly appearing reversal mutation will not establish in the population due to clonal interference. Subsequently, transmission bottlenecks might contribute to the fixation of the compensatory mutation, after which the reversal mutation has only a low or no selective advantage precluding its establishment in the population (52). Thus, in the MTB genetic architecture with widespread positive epistasis, compensatory variants might have a higher likelihood of being fixed compared to reversal mutations, even when the combination of resistance and compensatory mutation has a lower fitness than the reversal mutation. This evolved genetic architecture thus supports the fixation of compensatory mutations instead of reversal mutations.

Here we highlight that inherited indels were found to co-occur with substitutions. Although some of the co-occurring variants might be explained by genetic hitchhiking, the presence of co-occurring indels and substitutions in related gene functions or pathways supports that these variants interact epistatically. We thus conclude that MTB evolved a genetic architecture with widespread positive epistasis, where epistatic interactions between substitutions and indels contribute to the establishment of indels in the population.

Taken together, our results demonstrate the interplay between substitutions and indels in the evolution of biological functions that are essential for MTB infection and antibiotic resistance. We identified short and long indels that improve the resolution of outbreak phylogenies and that are crucial for the prediction of drug resistance in MTB strains. Especially for new hallmark drugs to treat multi-drug resistant MTB, such as bedaquiline and clofazimine, indels play a major role in collateral resistance towards both drugs (53,54). Thus, increasing knowledge on interactions between all variant types is paramount for our understanding of the fundamental evolutionary principles that govern the spread of antibiotic resistance and the associated compensatory mechanisms in MTB.

## Methods

### Sample collection and variants calling

We analyzed 353 multi-drug resistant MTB strains sampled longitudinally from the Central Asian Outbreak (MTB lineage 2, first referenced in (25)). Inclusion criteria were the presence of genetic markers defining the CAO clade (25). The strain collection was mainly assembled from a previously published collection derived from a drug resistance survey in Karakalpakstan, Uzbekistan (10) and from routine multi-drug resistance TB surveillance data from German patients (Table S1a) with 199 newly generated datasets and covering a sampling time from 1995 to 2015 (Table S1a). The closely related and fully drug-susceptible strain 49-02 (RefSeq: NZ_HG813240.1 version 11-MAR-2017) serves as the reference genome for variant calling. In addition, we performed the analysis for an outbreak of 64 fully drug-susceptible isolates (Table S1b) in the Hamburg region (MTB lineage 4, first referenced in (26)). Variants were inferred on the complete reference genome 7199-99 (RefSeq: NC_020089.1 version 19-MAY-2017).

We first trimmed the reads using trimmomatic v. 0.36 (55), with parameters `SLIDINGWINDOW:4:15 MINLEN:36 LEADING:3 TRAILING:3`. We mapped the trimmed reads to the outbreak reference genome using BWA MEM v0.7.16 (56), realigned around indels with GATK v3.8-0-ge9d806836 (57), and marked duplicates with PICARD v2.13.2. The median coverage ranges from 41 to 255. To detect SNPs and indels, we combined seven variant calling tools: GATK v3.8-0-ge9d806836 (57), FreeBayes v1.1.0-50-g61527c5 (58), Delly v0.7.7 (59), Pindel v0.2.5b9 (60), SvABA FH Version 134 (61), Scalpel v0.5.3 (62), and MindTheGap v2.0.2 (63). Tools were run and variants filtered according to tool-specific quality scores (Table S8) and we retained variants with a frequency over 75%.

Many indels occur in genomic regions of high GC content or that contain tandem repeats (37). The alignment of reads in these regions is therefore more difficult, and we expect to find higher uncertainty levels for indels compared to SNPs. Notably, in the case of MTB, this has led to the systematic exclusion of variants in PE and PPE genes (19). Here we implement two filtering steps, re-genotyping and back-genotyping, to obtain high-quality genomic variants.

### Re-genotyping

Since variants can be missed by variant calling, we performed re-genotyping to ascertain presences and absences of each variant in every sample. Re-genotyping determines whether the read alignment contains sufficient signal to support the variant. Additionally, we performed re-genotyping with a single tool per variant category. Therefore, the re-genotyped variant support values (i.e., the read support for the alternative allele) standardize the variant scores, allowing comparable assessment of variants from similar category. We used GATK to re-genotype SNPs and short indels and svtyper (64) for long deletions (Table S8). Recommended hard filtering was applied to the variants genotyped with GATK and all re-genotyped variants having a frequency over 75% were retained. Since no tool, to our knowledge, can be used to genotype long insertions, we used the breakpoints identified by MindTheGap as evidence of presence.

### Back-genotyping

To quantify the uncertainty associated with the variant calling, we implemented an additional layer of filtering for ambiguous signal. The idea behind the back-genotyping is to identify the “inverse signal” of a variant to confirm the absence of a variant (Fig S1). The approach consists of (i) generating multiple modified genomes of the outbreak reference that contain the detected variants, (ii) mapping the reads of each sample to each of the modified genomes, and (iii) genotyping the variant positions as for re-genotyping. For SNPs, we genotype the variants as they were introduced. For insertions, we genotype the putative deletions; for deletions, we genotype the putative insertions.

Variants in genomic regions that are difficult to align are expected to exhibit many uncertainties, i.e. contradicting results between genotyping and back-genotyping. Therefore, we filtered SNPs if they had more than five uncertainties, and the remaining variants if they had more than 20 uncertainties. The threshold has been determined in order to limit the incompatible variants in the final set of variants (Fig S2).

### Functionality assignments & enrichment tests

Since functions of MTB genes are determined based on experiments in H37Rv, we first retrieved homologs between the outbreak reference and H37Rv (NC_000962.3). For this, we performed a blast all-vs-all (65) between both sets of protein sequences and retrieved the significant hits (e-value < 1e-10). After computing the global identity between significant hits with the needle algorithm (66,67), we considered proteins as homologs if they shared a global identity higher than 30%. We then assigned the six essentiality categories described in (27) to the homologous proteins in the outbreak reference and grouped the gene categories into two main ones: essential genes are genes that are required for growth *in vitro*, and dispensable genes are genes that are not required for the growth *in vitro*. The latter category includes genes annotated as non-essential, conferring growth advantage, conferring growth defect, uncertain and containing an essential domain and genes without homolog in H37Rv.

A list of genes that were found to confer drug-resistance upon mutations in H37Rv are additionally used for annotation (Table S2). We identified the homologs in the outbreak reference and assigned the ABR-conferring or non-ABR-conferring categories accordingly.

### Phylogeny inference and evolutionary rate estimation

The back-genotyping allowed us to generate a presence-absence matrix, where uncertainties are represented as gaps. We estimated the phylogeny based on the presence-absence patterns of the final variants with iqtree v1.6.1 (68), using the GTR2+FO binary model, and the ultrafast bootstrap.

We displayed the tree using iTol v4.3.3 (69). The position of the root was estimated by the least-squares dating method implemented in LSD (70). Variants that are absent and present at least two times each in the presence-absence pattern contain information on the phylogenetic relationships, i.e. they are parsimony informative.

We used LSD v0.3beta (70) to estimate evolutionary rates. We followed Menardo et al. (2019) to determine the significance of the temporal signal (22). For each class of variants, we extracted their presence-absence sub-matrix, estimated the branch lengths on the pre-estimated phylogeny (iqtree option –pre), and performed the date-randomization test (DRT) (29). The calculated evolutionary rates are considered significant with three grades of significance. The stringent test assigns significance if the calculated confidence intervals of the rate do not overlap with the DRT confidence intervals. The intermediate test assigns significance if the calculated rates do not overlap with the DRT confidence intervals. The simple test assigns significance if the calculated rates do not overlap with the DRT rates (22). The hard limit of 1e-10 is imposed by LSD.

## Acknowledgements

We thank V. Mohr, F. Boysen and T. Ubben for excellent technical assistance for next generation sequencing of isolates. We thank Claudio Köser from the University of Cambridge for providing the list of antibiotic resistance-conferring genes in MTB. We thank Tobias Marschall from the Max-Planck Institute for Informatics Saarbrücken for discussions on variant calling and genotyping.

## Author contributions

AK, TD, SN, MG, MM, and TAK designed the study; MM, TAK, RD, FM, and SN collected the data; MG analyzed the data; MG, AK, and TD interpreted the results; MG and AK wrote the manuscript with contributions from all authors.

