## Supplementary figures for "Insertion and deletion evolution reflects antibiotics selection pressure in a *Mycobacterium tuberculosis* outbreak"

### Supplementary tables

**Table S1. Samples and accession numbers**. (a) Central Asian Outbreak. (b) Hamburg outbreak.

**Table S2. Antibiotic resistance-conferring genes.**

**Table S3. Number of variants in each category and raw numbers used for the enrichment analysis.** The essential category can be either essential (i.e., the gene is required for growth *in vitro*) or dispensable (i.e., the gene is not required for growth *in vitro*). "Antibiotic resistance" describes whether the gene has been found to confer antibiotic resistance upon substitutions ("Res", according to Table S2) or not ("NRes"). The p-value is the result of Fisher's exact test, with FDR-adjusted values.

**Table S4. Localization and description of the effect of parsimony informative indels.** The column "Parsimony score" gives the number of independent events required to explain the evolution of the variant on the phylogeny. Parsimony scores above 1 indicates incompatible variants. "Number of samples" describes the number of samples affected by the variant. Effect is either IS6110 insertion, homopolymer for an indel in a region with at least 3 repeated nucleotides, in-frame if the indel preserves the frame of the gene, or other for the remaining indels. The essential category can be either essential (i.e., the gene is required for growth *in vitro*) or dispensable (i.e., the gene is not required for growth *in vitro*). "Antibiotic resistance" describes whether the gene has been found to confer antibiotic resistance upon substitutions ("Res", according to Table S2) or not ("NRes"). "Comment" describes whether a gene is partially or completely deleted in the case of long deletions or the genomic distance of intergenic variants from the neighboring genes. Intergenic variants are split in two rows, where the first row describes the gene before the variant, and the second row describes the gene after the variant, in genomic coordinates. Of note, four of the incompatible short indels are found in homopolymer regions. In addition, four intergenic short indels that are incompatible likely stem from the same event, potentially involving a DNA translocation. The two remaining incompatible short indels constitute (i) an inframe short deletion in a PE gene and (ii) a 1bp insertion leading to a frameshift (and a modified protein sequence that is 28 amino acids longer), in a gene encoding a nitronate monooxygenase.

**Table S5. Description of incompatible SNPs.** The column "Effect" gives the effect of the SNPs on the resulting protein, i.e., non-synonymous (NS) or synonymous (S). "Parsimony score" gives the number of independent events required to explain the evolution of the variant on the phylogeny. "Number of samples" describes the number of samples where the variant is detected. The essential category can be either essential (i.e., the gene is required for growth *in vitro*) or dispensable (i.e., the gene is not required for growth *in vitro*). "Antibiotic resistance" describes whether the gene has been found to confer antibiotic resistance upon substitutions ("Res", according to Table S2) or not ("NRes"). "Comment" describes the genomic distance of intergenic variants from the neighboring genes. Intergenic variants are split in two rows, where the first row describes the gene before the variant, and the second row describes the gene after the variant, in genomic coordinates.

**Table S6. Summary of genes affected by convergent evolution due to indels.** "Parsimony informative" states whether the variant considered is parsimony informative (Yes) or not (No). "Parsimony score" gives the number of independent events required to explain the evolution of the variant on the phylogeny. "SNPs found elsewhere" indicates the number of SNPs found in the gene considered. The essential category can be either essential (i.e., the gene is required for growth *in vitro*) or dispensable (i.e., the gene is not required for growth *in vitro*). "Antibiotic resistance" describes whether the gene has been found to confer antibiotic resistance upon substitutions ("Res", according to Table S2) or not ("NRes").

**Table S7. dN/dS analysis for genes having at least two SNPs.** Genes are sorted by the number of SNPs they contain. “Antibiotic resistance” describes whether the gene has been found to confer antibiotic resistance upon substitutions (“Res”, according to Table S2) or not (“NRes”). The essential category can be either essential (i.e., the gene is required for growth *in vitro*) or dispensable (i.e., the gene is not required for growth *in vitro*).

**Table S8. Command lines and parameters for variants calling and regenotyping.**

### Supplementary figures

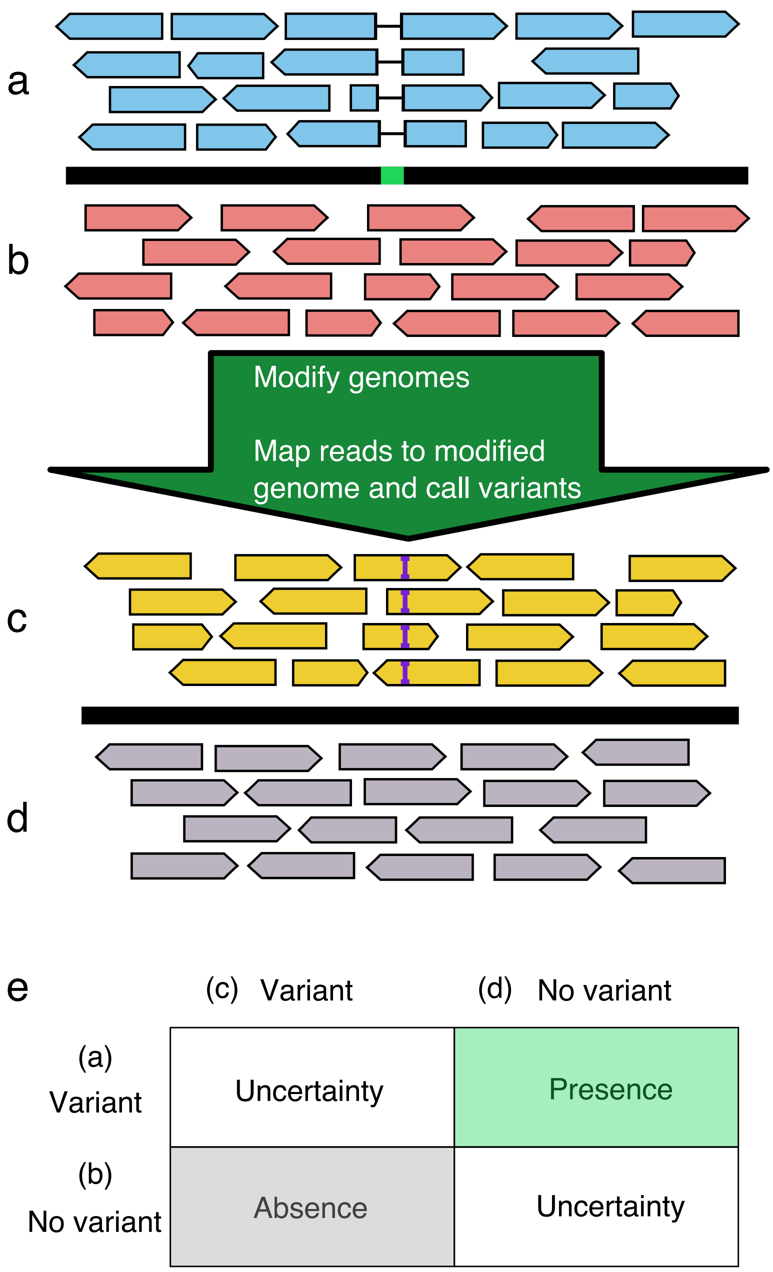


**Fig S1. Schematic summary of the back-genotyping.** The example shows the calling and back-genotyping of a small deletion. (a)-(b) We identify a short deletion in one sample (blue). We include this deletion in the modified genome and map all the samples, and we genotype variants. (c)-(d) We either detect an insertion (yellow sample), if the base pairs that are removed in the modified genome are present in the reads, or nothing if the deleted portion is not present in the reads (gray sample). (e) The decision table summarize the decision on the variants for each sample. If a variant is detected in the first phase and nothing is detected in the second phase, the variant is considered present. Alternatively, if the variant was not detected in the first phase but the inverse variant is detected in the second phase, the variant is considered absent. For the remaining cases, the variant is considered uncertain. These decisions are summarized in a presence-absence matrix, with presence (1), absence (0) and uncertainty (gap, "-").

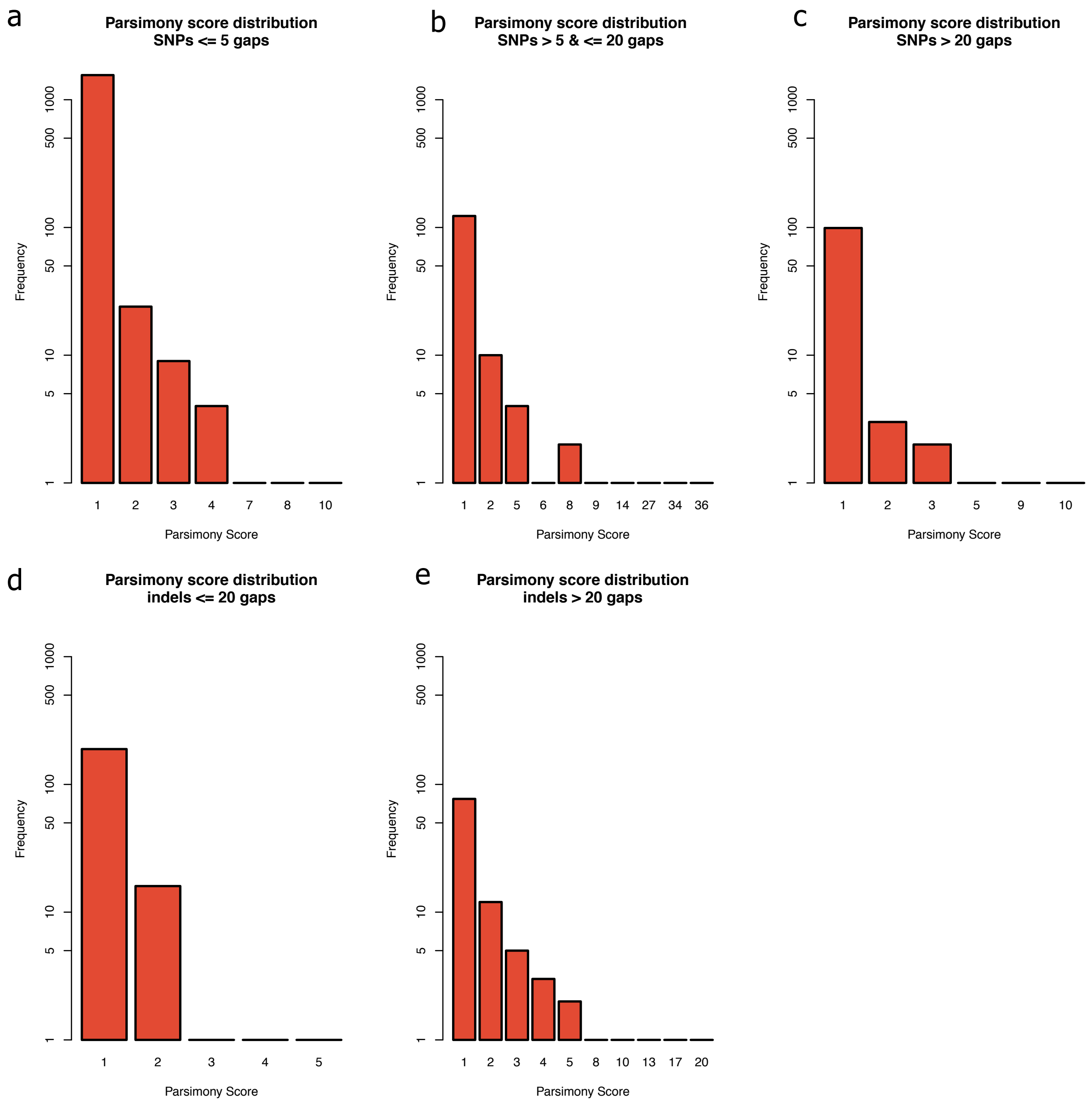


**Fig S2. Gap counts and parsimony scores for SNPs and indels.** (a) SNP having less than five gaps, (b) SNP having between six and 20 gaps. (c) SNPs having more than 20 gaps, (d) indels having at most 20 gaps, (e) indels having more than 20 gaps. The parsimony scores of SNPs between 5 and 20 gaps are high (up to 36), therefore we excluded SNPs having more than 5 gaps. Indels exhibit rather low parsimony scores (1 to 5) for variants exhibiting at most 20 gaps. Variants having more than 20 gaps have higher parsimony scores (up to 20), therefore the threshold has been set at 20 gaps for including indels.

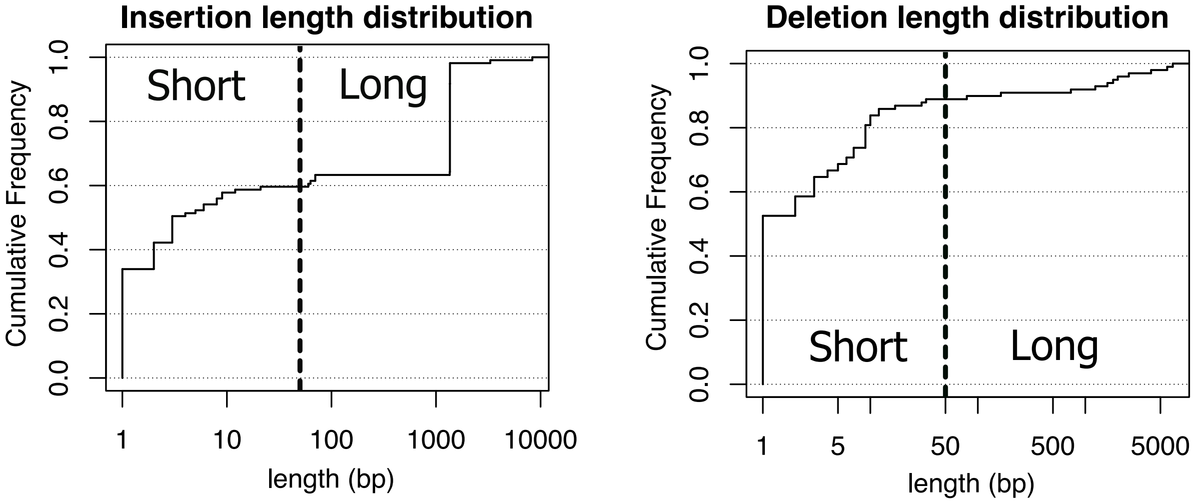


**Fig S3. Cumulative length distribution of 208 indels.** 65 of 44 insertions are short (59.6 %) and 88 of 99 deletions are short (88.9%).

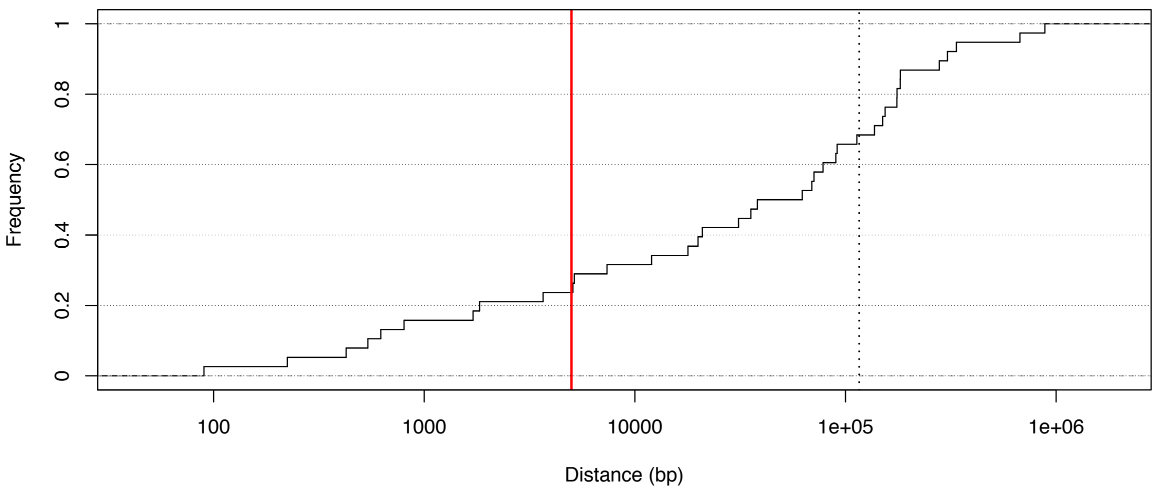


**Fig S4. Cumulative distribution of the distance between neighboring IS6110 elements.** An IS6110 insertional hotspot is defined by at least two insertions in two different samples, where the distance between neighboring insertion sites is at most 5000 bp. This cutoff was chosen since it includes 10 of 38 neighboring pairs (26.3%, red line). The grey dotted line shows the expected distance between neighboring IS if they are uniformly distributed along the genome (116,115 bp). In our data, we could classify 17 of the 38 IS insertions (44.7%) in seven hotspots. Two of the hotspots have already been reported and they correspond to insertions into the DnaA-DnaE intergenic region and the phospholipase C region. The five remaining hotspots are located in genomic regions consisting mainly of hypothetical proteins.

a

| | SNPs | Insertions | | Deletions | | Total |
|---|---|---|---|---|---|---|
| | | Short | Long | Short | Long | |
| **In Gene** | 82 (85.4%) | 0 | 0 | 10 (83.3%) | 0 | **92** |
| **Intergenic** | 10 (10.4%) | 0 | 0 | 0 | 0 | **10** |
| **In pseudogene** | 4 (4.16%) | 0 | 2 (100%) | 2 (16.7%) | 2 (100%) | **10** |
| **Total** | **96** | **0** | **2** | **12** | **2** | **112** |
| **Parsimony informative** | 34 (35.4%) | 0 | 1 (50%) | 6 (50%) | 1 (50%) | **42** |
| **Compatible** | 34 (100%) | 0 | 1 (100%) | 5 (83.3%) | 1 (100%) | **41** |

b

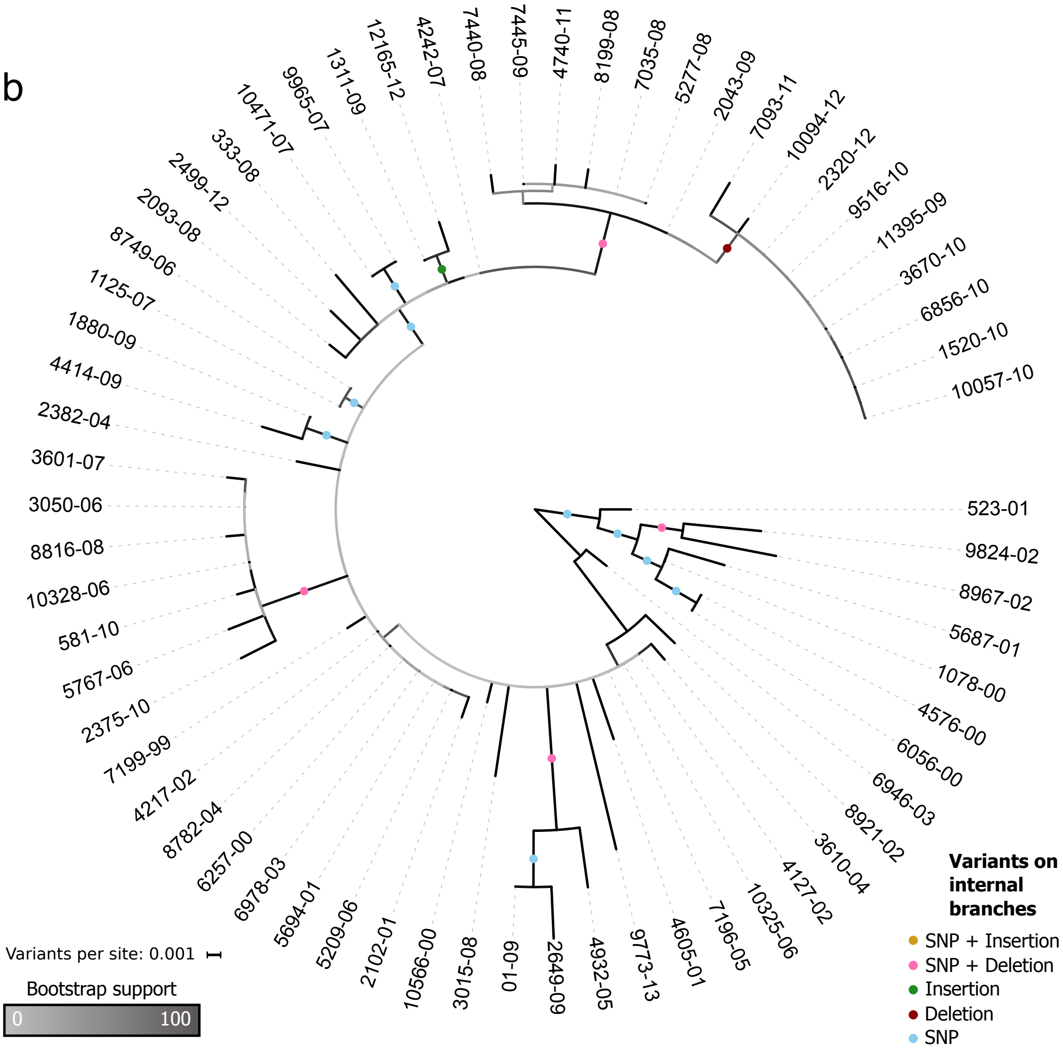


c

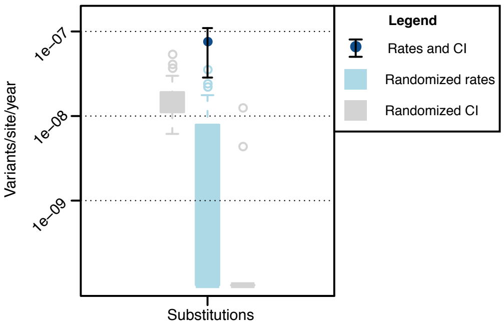

**Fig S5. Genomic localization, phylogenetic tree, evolutionary rates and enrichment analysis for 64 strains sampled from the "Hamburg" outbreak.** The Hamburg outbreak is a fully-sensitive MTB strain of lineage 4 sampled in the Hamburg and Schleswig-Holstein region between 1998 and 2013 (Table S1b; Roetzer et al., 2013). We performed the same analysis as the CAO for calling variants, including regenotyping and back-genotyping. No variants were found in ABR-conferring genes and the enrichment analysis on the essential category did not show any significant results. (a) Summary and genomic localization of detected variants. Percentages are calculated based on the total number of variants in each variant class, except for the compatible variants, where the percentage is calculated based on the number of parsimony informative variants in each variant class. (b) Phylogenetic tree of the Hamburg outbreak, inferred from the presence-absence patterns of 112 detected variants and rooted by the temporal root estimated with LSD. Two branches are refined by single indels. One is an IS6110 insertion inferred in a pseudogene annotated as a Fic protein, that groups two samples. The second is a 45bp deletion in a PE gene that groups nine samples. (c) Substitution rate and 95% confidence intervals (CI) estimated with LSD. Substitutions have temporal signal accorded by the intermediate test for temporal signal, with an estimated substitution rate of 7.51e-8 [2.85e-8 – 11.0e-8] substitutions/site/year, where the confidence interval (CI) includes the previous estimate of 1e-7 substitutions/site/year (Roetzer et al., 2013). Due to the limited number of events, there is not sufficient temporal signal to estimate evolutionary rates for indels.
